# Relevance of screening for Chagas and viral hepatitis in Bolivian migrants

**DOI:** 10.1101/775890

**Authors:** Irene Losada Galván, Giuseppe Gariup, Aina Casellas, Carme Subirà, Alex Almuedo-Riera, Daniel Camprubí, Natalia Rodríguez-Valero, Joaquim Gascón, Jose Muñoz, María Jesús Pinazo

## Abstract

**Objectives:** given the scarcity of data regarding prevalence of various infectious diseases in Latin-American countries, our study aims to assess the burden of *T.cruzi, S.stercoralis*, HIV and viral hepatitis in Latin-American migrants, with a focus on Bolivian migrants.

**Methods:** we performed a retrospective observational study of 565 screening evaluations on adults (≥ 18 years) carried out at our referral International Healthcare service in Barcelona. We reviewed structured clinical records and microbiological results of patients attended between February 2012 and April 2015.

**Results:** the median 35 years old and 74% were women. Bolivian origin accounted for 87% of the screened population. We found a 48% prevalence of T.cruzi, 16% of S.stercoralis, 0.2% of HIV, 92% of HAV, 0.2% HBV and 0.2% HCV.

**Conclusions:** these results support the relevance of the screening of *T. cruzi* and *S. stercoralis* in Bolivian migrants, but challenge the pertinence of systematic screening of HBV in this population.

**Author summary:** In response to the challenge of detecting diseases not previously present in host countries, screening programs have been implemented for migrants based on the probability of having certain diseases depending on their country of origin and / or migratory route. This increased risk is very clearly established in some cases such as *Trypanosoma cruzi* infection (the cause of Chagas disease) in people from Latin America; especially from Bolivia. In recent years screening recommendations for *Strongyloides stercoralis* in this population was proven necessary. Current recommendations regarding systematic screening for hepatitis B establish the relevance of screening based on the probability of the disease in the 2% population of origin. Since there are no reliable and up to date data regarding prevalence of hepatitis B virus in Bolivia, we aimed to analyze data available for migrants from Bolivia in Spain.

Our results support the importance of screening for *T. cruzi* and *S.stercoralis* in patients from Bolivia. However, our data show a much lower prevalence of this hepatitis B virus (0.2%) than the 2% threshold that would justify systematic screening, so we question the relevance of screening for hepatitis B virus in this population in the absence of other risk factors.

## Introduction

The increase in global migratory movements in recent decades implies new challenges for health professionals and policy makers, especially regarding the need to diagnose and treat previously non-endemic pathologies in the host countries. One of the diseases that best illustrates this challenge is Chagas disease, caused by chronic infection by *Trypanosoma cruzi* parasite.

It is estimated that 6-7 million people worldwide are infected with *T.cruzi*, mainly in Latin America (LA), where it causes more than 10000 deaths per year(1). The first reports of Chagas disease in Europe were published in the early 1980s, with a marked increase around 2000, along with the rising flow of migrants from LA to Europe(2–4). From 2007 onwards, a number of initiatives both at national and international levels have been implemented in order to increase awareness and provide better care to potentially affected populations (5). These aim for better control vertical transmission (congenital, from mother to fetus)(6) and transmission by infected blood and organ donors(7,8). Such initiatives include screening programs, which are often the first contact of the migrant with the health system in host countries. Thus, they represent an opportunity for a thorough medical examination and complementary tests. The indication of performing these diagnostic tests derives from the estimated prevalence for each risk group, using the country of origin in the absence of other known risk factors (9,10).

The relevance of conducting Chagas disease screening in Latin American migrants is based on epidemiological data from the countries of origin as well as data from recipient countries: WHO estimates a global prevalence of 1% in LA countries(11), with great variations within the different countries in the region (highest prevalence in Bolivia 6.1% and Paraguay 2.1%). Data from non-endemic areas shows similar results: a meta-analysis(12) on prevalence of Chagas disease in Europe estimated a prevalence of 4.2%, also with important variations according to country of origin, with the highest prevalence among Bolivian immigrants (18%). In addition, there are studies in favor of the cost-effectiveness of screening for Chagas disease in this context(13,14).

Another well-established indication for screening would be *Strongyloides stercoralis*, with an estimated prevalence of 370 million worldwide (15) and regional LA country-specific prevalence ranging from 1 to 73%(16,17). *S. stercoralis* chronic infection is usually asymptomatic, but it can develop into a severe and highly lethal disease in the context of immunosuppression(18). The association of *T.cruzi* infection with a two-fold increase in the odds of strongyloidiasis (19) support this combined strategy of screening (*T.cruzi - S.stercoralis*) in LA migrants.

WHO estimates that in 2015, 257 million persons were living with chronic Hepatitis B Virus (HBV) infection in the world, 68% in African and Western Pacific regions(20). Chronic HBV infection is usually asymptomatic, but 20-30% of patients with chronic VHB infection will develop complications (including liver cirrhosis and hepatocellular carcinoma). Testing for chronic HBV infection meets established public health screening criteria(21). In the absence of specific risk factors such as blood transfusion recipients before 1991, injectable drug users, men who have sex with men; a geographic-based screening is performed based on the HBV prevalence of the countries of origin. Before 2008 the HBV prevalence threshold was established at 8% (high endemicity countries), that was then lowered to a 2% prevalence (medium endemicity) in the updated recommendations of the CDC(22). Ever since there is a broad consensus on the relevance and cost-effectiveness of screening for hepatitis B in people from countries with prevalence greater than 2%(23). However this figure is currently under discussion: a study performed in the United States argued that it would be cost-effective screening in populations with prevalence as low as 0,3%(24). Both European(25–27) and American (28–30) guidelines recommend screening in migrants coming from countries with a HBV prevalence of 2% or higher. However, the discussion of the screening threshold becomes futile when epidemiological studies on which the recommendations are based by each country are scarce, not updated and often conflicting. The WHO estimates an overall prevalence in the region of the Americas between 0.4 and 1.6%(31), with 7-12 million Latin Americans carrying HBV chronic infection(32). In the case of Bolivia, there is conflicting evidence with regard to its HVB infection prevalence. A meta-analysis(33) showed a prevalence of between 0.1 and 6%, but its baseline data oscillate between 1987 and 2008, with a total sample of 1930 patients.

Another systematic review (34) estimates a prevalence of HBV in Bolivia of 0.44% based on 4 studies (1357 patients).

The scenario of chronic infection with hepatitis C virus (HCV) is very different. The prevalence of this disease is higher in Spain (1.7%) compared to the prevalence in Latin American countries (1.1%-1.3%, Bolivia 0.9%) (27,35). Therefore, given the low prevalence in LA countries, screening indication would be given in any case by risk factors different from the country of origin. However, some authors argue that HCV screening should be performed anytime HBV screening is indicated for another reason.

According to 2016 ECDC epidemiological assessment(36), estimates of total number of migrants infected with HCV or HBV might be an overestimation, since prevalence if often lower in migrants compared to the prevalence of the country of origin.

Data on serologic prevalence of viral hepatitis in LA migrants in Spain is scarce and often conflicting with reports from other countries in Europe. A report from a Tropical Medicine Centre in Madrid (37) found 1.6% chronic HBV infection in LA migrants. These results are similar to a 1.6% and 1.2% chronic HBV infection from a referral center (38) and a primary care study (39) both in Barcelona; as opposed to a 0.6% overall prevalence found on another study in the UK(40).

Considering the scarcity of data of the previously stated entities in LA migrants, our study aims to assess the burden of these infectious diseases in Latin-American migrants attending a referral International Healthcare service in Barcelona, with a focus on Bolivian migrants. We intend to contribute our data to those of other migrant cohorts in high income countries, given that they are a useful source of information on the prevalence of various infectious diseases when reliable data from the countries of origin are not available. We evaluate the number of chronic infections in the target population in comparison with that of their countries of origin in order to assess the pertinence of the systematic screening in this population.

## Methods

This retrospective observational study was performed at the International Health department of the Hospital Clinic of Barcelona. We reviewed medical records from all patients at risk of *T.cruzi* infection attending our unit between February 2012 and April 2015.

In our clinic we perform screening evaluations on adults (age 18 or older) who come from Latin American Countries. They come to the clinic either spontaneously, counseled by friends or relatives or referred by their physician. The first visit consists of an interview in which epidemiological information is collected as well as relevant medical history. A physical examination is performed and diagnostic tests are requested according to current screening guidelines.

Epidemiological data include age, sex, country of origin, year of arrival in Spain, risks factors for *T. cruzi* acquisition (rural area, adobe housing, contact with triatomine vector, blood transfusions, mother affected with Chagas disease) and risk factors for hepatitis (blood transfusions, unprotected sexual relationships). The usual screening workup includes a blood cell count, general biochemistry, serologies for *T. cruzi, S. stercoralis*, HIV, HBV and HCV. HAV was tested according to physician preference. Stool samples are collected for parasitological examination.

Screening of Chagas was performed with a chemiluminescent microparticle immunoassay (ARCHITECT Chagas^®^, Abbott)(41). A positive result was confirmed with a conventional ELISA with recombinant antigens (CHAGAS ELISA IgG+IgM, Vircell)(42) as per WHO recommendations for diagnosis(43). *S. stercoralis* infection was diagnosed by direct visualization of ova in stool or a positive one-step sandwich-format immunoassay for the qualitative detection of IgG-class antibodies to *Strongyloides stercoralis* antigen (Strongyloides ELISA, SciMedx)(44). Detection of hepatitis B virus surface antigen (HBsAgII, Advia Centaur) (45), antibody against hepatitis B virus core (HBc Total, Advia Centaur) (46) and surface antigen (antiHBs 2, Advia Centaur)(47) were used to assess HBV status. Active infection was considered if HBsAg was positive; vaccination if HBsAg was negative, HBsAb positive and HBcAb negative, and cured infection if HBsAg is negative and HBsAb and HBcAb positive. Accurate classification was not always possible since not all the individuals in the cohort had a complete serology. Antibodies against hepatitis A (HAV Total, Advia Centaur)(48) and C (HCV, Advia Centaur)(49) viruses were used to detect passed HAV infection and to screen for chronic HCV infection. Screening of HIV was made by a chemiluminometric immunoassay of antigen binding microparticles that is used to detect antibodies against human immunodeficiency virus type 1, including subtype O, and / or type 2 (HIV 1 / O / 2 Enhanced Assay, Avia Centaur)(50).

Data were presented as frequencies and median (interquartile range, IQR) for discrete and continuous variables, respectively. Proportions were compared using Chi-squared test or Fisher’s exact test if the application conditions of the former where not met. Medians were compared between groups using Wilcoxon Rank Sum test. Significance was set at 0.05. The analysis was carried out using Stata 15 (StataCorp. 2017)(51).

This study was approved by the ethics committee for medical research of Hospital Clinic Barcelona with reference number HCB/2018/0521. All analyzed data were previously anonymized.

## Results

Over the study period, 565 individuals were screened for *T. cruzi* and other infectious diseases. The median (IQR) age was 35 (29 - 42) years old and 74% were women. Median (IQR) time elapsed from their arrival to the country and this screening was eight (7–10) years. Demographic characteristics and presence of risk factors for *T. cruzi* infection of the study population compared by *T. cruzi* results are shown in Table 1. Four hundred and ninety-five Bolivian patients were screened, accounting for 87% of the screened population. The remaining 71 individuals came from other Latin American countries, with Argentina as the next most numerous country of origin (21, 4%).

**Table 1.**
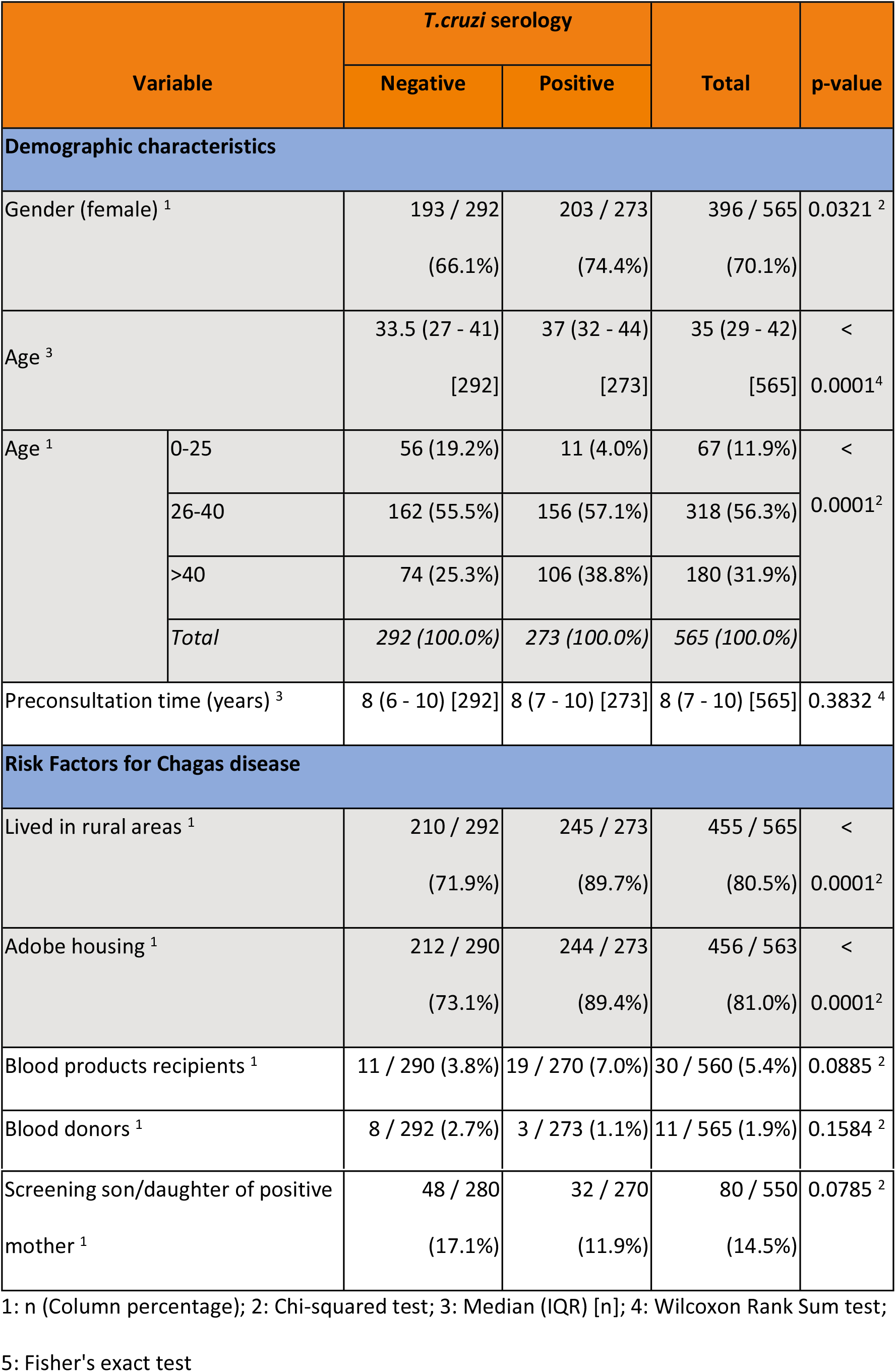
Demographic characteristics and *T. cruzi* risk factors

### Chagas disease

*T. cruzi* was positive in 273 participants (48%) of the screened individuals. Among them, there was a greater presence of known risk factors for *T. cruzi* infection as residence in rural area (90%, p <0.0001), adobe housing (89%, p <0.0001), but no differences were found regarding the receipt of blood products. Out of 80 patients whose mothers had confirmed positive serology for *T. cruzi*, 32 (40%; 95%CI: 29-52%) were also positive for *T. cruzi* but without significative difference.

We also compared *T.cruzi* positivity in relation to other infectious diseases, with results shown in Tables 2 and 3.

**Table 2.**
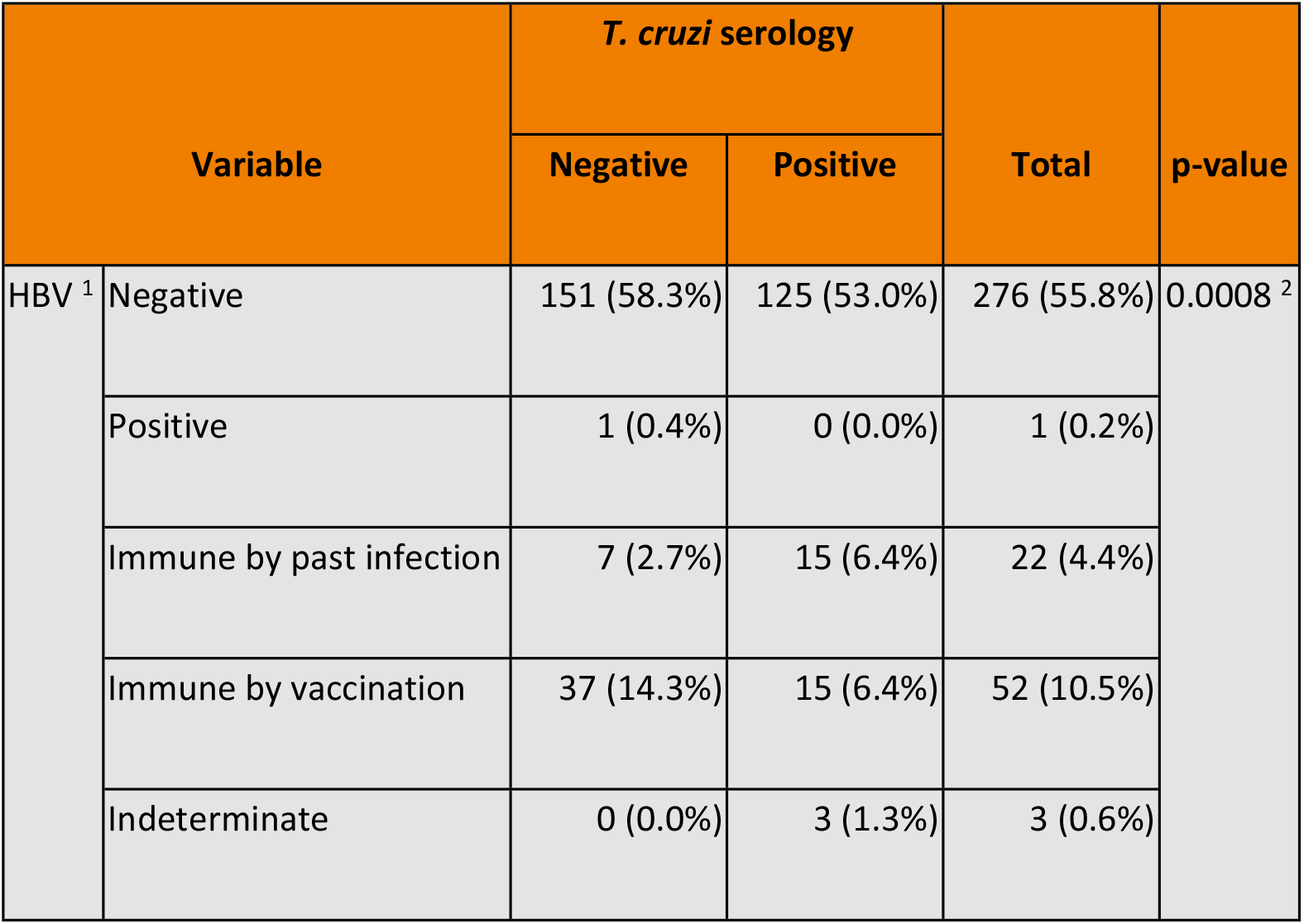
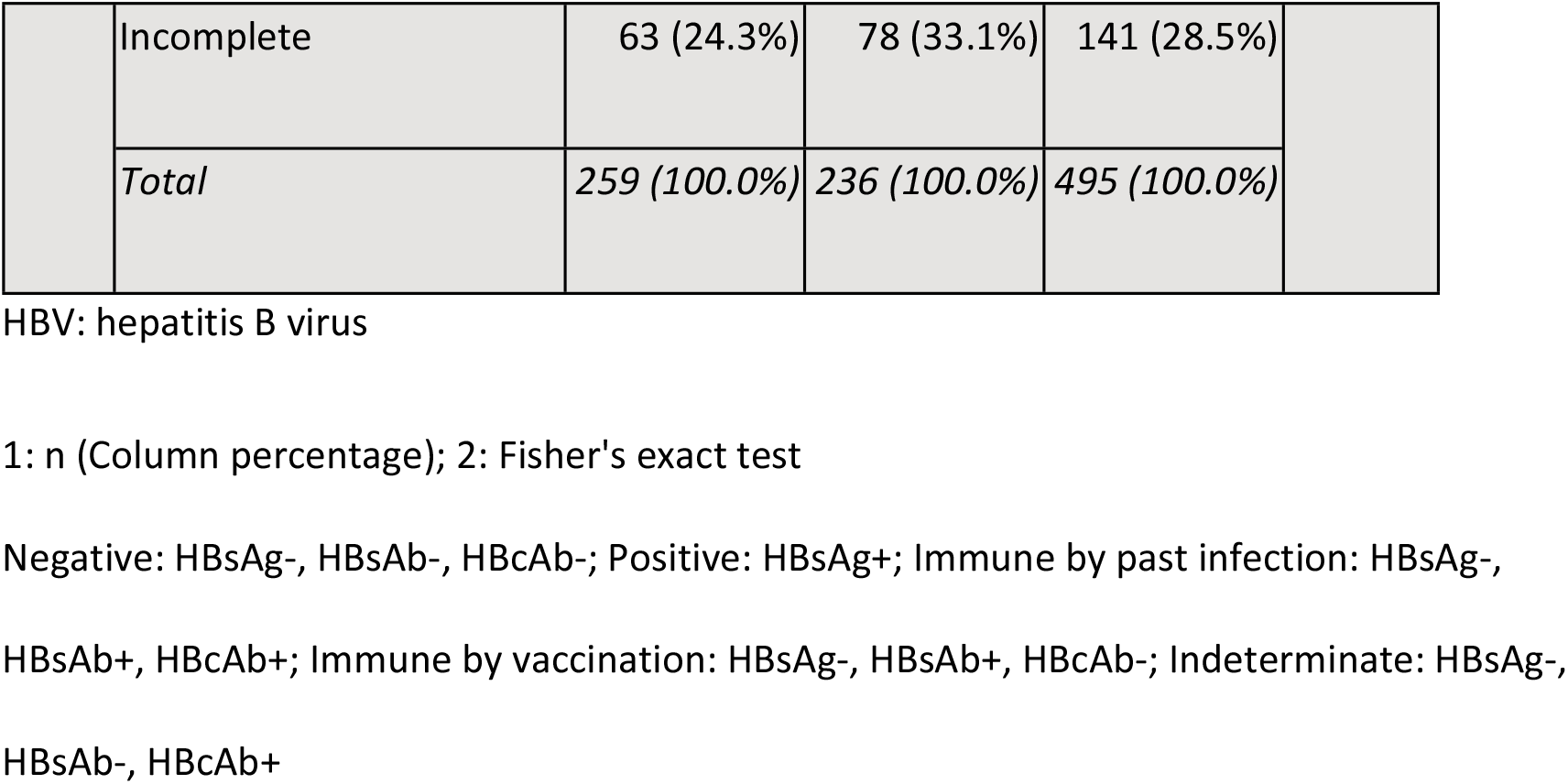
Hepatitis B virus screening by *T. cruzi* results

| Variable |  | *T. cruzi* serology |  | Total | p-value |
| --- | --- | --- | --- | --- | --- |
|  |  | Negative | Positive |  |  |
| HBV <sup>1</sup> | Negative | 151 (58.3%) | 125 (53.0%) | 276 (55.8%) | 0.0008 <sup>2</sup> |
|  | Positive | 1 (0.4%) | 0 (0.0%) | 1 (0.2%) |  |
|  | Immune by past infection | 7 (2.7%) | 15 (6.4%) | 22 (4.4%) |  |
|  | Immune by vaccination | 37 (14.3%) | 15 (6.4%) | 52 (10.5%) |  |
|  | Indeterminate | 0 (0.0%) | 3 (1.3%) | 3 (0.6%) |  |
|  | Incomplete | 63 (24.3%) | 78 (33.1%) | 141 (28.5%) |  |
|  | *Total* | *259 (100.0%)* | *236 (100.0%)* | *495 (100.0%)* |  |

HBV: hepatitis B virus
1: n (Column percentage); 2: Fisher's exact test
Negative: HBsAg-, HBsAb-, HBcAb-; Positive: HBsAg+; Immune by past infection: HBsAg-,
HBsAb+, HBcAb+; Immune by vaccination: HBsAg-, HBsAb+, HBcAb-; Indeterminate: HBsAg-,
HBsAb-, HBcAb+

**Table 3.** Other infectious diseases by *T. cruzi* results

| Variable |  | T. cruzi serology |  | Total | p-value |
| --- | --- | --- | --- | --- | --- |
|  |  | Negative | Positive |  |  |
| Screening for other infectious diseases |  |  |  |  |  |
| HIV | Negative | 270 (99.6%) | 253 (100.0%) | 523 (99.8%) | 1.0000 <sup>3</sup> |
|  | Positive | 1 (0.4%) | 0 (0.0%) | 1 (0.2%) |  |
|  | Total | 271 (100.0%) | 253 (100.0%) | 524 (100.0%) |  |
| HAV IgG <sup>1</sup> | Negative | 15 (11.5%) | 5 (3.9%) | 20 (7.8%) | 0.0229 <sup>2</sup> |
|  | Positive | 115 (88.5%) | 122 (96.1%) | 237 (92.2%) |  |
|  | Total | 130 (100.0%) | 127 (100.0%) | 257 (100.0%) |  |
| HCV <sup>1</sup> | Negative | 234 (100.0%) | 223 (99.6%) | 457 (99.8%) | 0.4891 <sup>3</sup> |
|  | Positive | 0 (0.0%) | 1 (0.4%) | 1 (0.2%) |  |
|  | *Total* | *234 (100.0%)* | *224 (100.0%)* | *458 (100.0%)* |  |
| Strongyloides stercoralis <sup>1</sup> | Negative | 178 (87.7%) | 152 (76.8%) | 330 (82.3%) | 0.0123 <sup>5</sup> |
|  | Positive | 23 (11.3%) | 42 (21.2%) | 65 (16.2%) |  |
|  | Indeterminate | 2 (1.0%) | 4 (2.0%) | 6 (1.5%) |  |
|  | *Total* | *203 (100.0%)* | *198 (100.0%)* | *401 (100.0%)* |  |

HIV: Human immunodeficiency virus; HAV: hepatitis A virus; HBV: hepatitis B virus; HCV:

### HIV

Serologic test for HIV was performed in 93% of the study population, with one positive result on a man who had sex with men, without other risk factors.

### Hepatitis A virus

A serologic test for HAV was performed in 45% of the study population. A positive HAV was significantly more frequent among individuals who were also positive for *T.cruzi*, with an overall prevalence of 92%. (Table 3)

### Hepatitis B virus

A serologic test for HBV was performed in 88% (495) of the study population. However, there was insufficient information for HBV classification on 141 individuals (28%). Among those with a complete serology, there was a 7% prevalence of positive serology for HBV (either by chronic or immune by past infection) –see Table 2-, with one case (0.2%) of chronic HBV. This patient had been previously diagnosed on a routine test in Bolivia. She denied risky sexual practices, transfusions or parenteral drug use. However, she reported several accidental punctures when he practiced dentistry in Bolivia.

The remaining 93% had a negative serology for HBV or were immune by vaccination. The prevalence of chronic/past infection was significantly higher among individuals who were also positive for *T. cruzi*. Table 4 shows demographic characteristics and microbiological results of Bolivian patients by HBV.

**Table 4.** Results from Bolivian migrants with respect to HBV results

| Variable |  | HBV |  | Total |
| --- | --- | --- | --- | --- |
|  |  | Negative or Immune by vaccination | Positive or Immune by past infection |  |
| Gender (female) <sup>1</sup> |  | 195 / 287 (67.9%) | 13 / 22 (59.1%) | 208 / 309 (67.3%) |
| Age <sup>2</sup> |  | 36 (28 - 43) [287] | 36 (33 - 46) [22] | 36 (29 - 43) [309] |
| Age <sup>1</sup> | 36 (12.5%) | 0 (0.0%) | 36 (11.7%) | 36 (12%) |
|  | 158 (55.1%) | 16 (72.7%) | 174 (56.3%) | 174 (56%) |
|  | 93 (32.4%) | 6 (27.3%) | 99 (32.0%) | 99 (32%) |
|  | 287 (100.0%) | 22 (100.0%) | 309 (100.0%) | 309 (100%) |
| Preconsultation time (years) <sup>2</sup> |  | 8 (7 - 9) [287] | 8 (7 - 10) [22] | 8 (7 - 9) [309] |
| Screening for other infectious diseases |  |  |  |  |
| HIV <sup>1</sup> | Negative | 284 (100.0%) | 22 (100.0%) | 306 (100.0%) |
| HAV IgG <sup>1</sup> | Negative | 9 (6.8%) | 3 (27.3%) | 12 (8.3%) |
|  | Positive | 124 (93.2%) | 8 (72.7%) | 132 (91.7%) |
|  | Total | 133 (100.0%) | 11 (100.0%) | 144 (100.0%) |
| HCV <sup>1</sup> | Negative | 257 (99.6%) | 20 (100.0%) | 277 (99.6%) |
|  | Positive | 1 (0.4%) | 0 (0.0%) | 1 (0.4%) |
|  | *Total* | *258 (100.0%)* | *20 (100.0%)* | *278 (100.0%)* |
| *Strongyloides stercoralis*<br>serology <sup>1</sup> | Negative | 146 (50.9%) | 11 (50.0%) | 157 (50.8%) |
|  | Positive | 30 (10.5%) | 5 (22.7%) | 35 (11.3%) |
|  | Indeterminate | 5 (1.7%) | 0 (0.0%) | 5 (1.6%) |
|  | Missing | 106 (36.9%) | 6 (27.3%) | 112 (36.2%) |
|  | *Total* | *287 (100.0%)* | *22 (100.0%)* | *309 (100.0%)* |

HIV: Human immunodeficiency virus; HAV: hepatitis A virus; HCV: hepatitis C virus.
1: n (Column percentage); 2: Median (IQR) [n]

### Hepatitis C virus

A serologic test for HCV was performed in 81% of the study population. There was one positive result from a patient who was also positive for *T. cruzi*, but negative for HBV and HIV. This patient was initially screened for HCV in 2014 with a negative result. During follow-up, an elevation of transaminases advised a new a serological study, which was positive for HCV. She had not traveled since 2014; so acute HCV infection (genotype 1a) acquired in Spain between February and March 2016 was suspected. Her elastography showed no fibrosis and was successfully treated with Sofosbuvir/Ledipasvir in 2017 (viral load undetectable since then).

### Strongyloides stercoralis

A serologic test and/or stool sample for parasite examination was obtained in 71% of the study population, with an overall prevalence of 16%. This prevalence of *Strongyloides stercoralis* infection was significantly higher in patients who were also positive for *T. cruzi* (Table 3).

## Discussion

The aim of this study was to describe the prevalence of different infectious diseases in Bolivian and other Latin-American migrants attending a referral International Health service in order to establish more accurate screening protocols according to international recommendations. First we should acknowledge that the vast majority of the screened individuals were from Bolivia, and coming from several areas with a high migratory tradition (52). This limits the generalization of the results to the Latin American community as a whole, but provides with a robust source of information about the Bolivian community in our context. Hence, most of our analysis will apply only to Bolivian migrants.

Another relevant finding is a prevalence for ***T. cruzi*** infection as high as 48% of the screened population. This is probably due to various reasons. First, the majority of the screened population was from highly endemic areas in Bolivia (with an expected prevalence of 18%)(12). Secondly, some patients already know about their diagnosis before coming to our clinic. Moreover, patients with a positive result for T. *cruzi* might be more prone to follow the general advice of inviting their relatives to the screening program. Thus, this unusually high prevalence of Chagas disease might be influenced by community and family clusters that share risk factors for the disease. These risk factors include having lived in a house made of adobe and having lived in rural areas, which are both significant risk factors for *T. cruzi* in this study population. Interestingly, in our cohort, having received blood products was not statistically associated with a higher risk of *T. cruzi* infection. This might be due to the relative small number of patients who did receive some blood product in their countries of origin, but it might also be an early outcome of the blood bank controls that were implemented in Bolivia (43,54). In light of these results and others in other non-endemic countries (3,12), we cannot stress enough the need for standardized screening programs for *T. cruzi* in Latin American migrants. An early *T.cruzi* diagnosis allows the individual evaluation and adequacy of the support treatment according to the clinical stage. In addition, it enables the evaluation of the indication of antiparasitic treatment (55,56) in order to reduce the likelihood of progression of the disease as well as being an instrument for interrupting transmission, especially in non-endemic areas through the treatment of women of childbearing age (57–59).

In agreement with previous work (19), we found a high prevalence of **Strongyloidiasis** and an association with *T. cruzi* infection. This further supports maintaining a combined screening strategy of these two pathogens.

The observed high **hepatitis A virus** prevalence and its association with *T.cruzi* infection might be due to shared risk factors in terms of low socio economic status and deficient hygienic conditions, although the most frequent means of transmission are essentially different for these two diseases. However, this observation could be conditioned by the fact of having HAV data only of 45.5% of the patients.

Our results show very low chronic **hepatitis B virus** prevalence (0.2%), although up to 7% of the screened population was immune by past infection. This prevalence of chronic HBV would not justify a systematic screening program in these patients. The most accepted threshold is 2%(25–30), well above the most ambitious estimates which recommend screening in communities with a prevalence of 0.3% (24). However, it is possible that the selection of this cohort of patients is not random within Bolivia and that we are facing a healthy migrant bias. Moreover, the fact that patients are screened for HBV a median of 8 years after they arrive, further jeopardizes the probability of diagnosing acute infections. Acute infections acquired prior to departure or in the host country might be wrongly classified due to this long period between arrival and HBV screening. Moreover, no complete serological information was available for almost one third of patients, which constitutes one of the main limitations of the study. There are no updated data on prevalence of HBV or vaccination coverage since its introduction in 2000 (60). It would be necessary to have up-to-date and robust evidence on the epidemiology of HBV in Bolivia in order to improve the targeted screening of this disease in the absence of other risk factors for HBV. In any case, this scenario of young and healthy patients is common in countries that host migrant population; and therefore it is the basis on which estimates of the necessary screening can be made, in the absence of reliable epidemiological data from the countries of origin.

Due to the negative gradient of the prevalence of **hepatitis C virus** in Spain with respect to Bolivia, the systematic screening of this disease in the absence of other risk factors would not be granted. However, as some authors recommend, HCV serology is sometimes performed when requesting other serologies, such as HBV or HIV in young people who are sexually active due to the risk of acquisition in the host country. Our results confirm the relevance of this strategy in both the prevalence of HCV and HIV.

## Conclusions

This work supports the relevance of the screening of *T.cruzi* and strongyloides in people from Bolivia. Hepatitis A virus is endemic and the vast majority of people have positive serology due to past infection, therefore screening is not recommended. According to available evidence, the systematic screening of HCV and HIV is not recommended, with the exception of making at least one determination in sexually active persons. On the other hand, this work questions the relevance of systematic HBV screening in Bolivian migrants given its low prevalence. Epidemiological studies are urgently needed to clarify the situation of HBV in Bolivia in order to direct efforts both inside and outside its borders.

## Funding

The team is supported by the Agencia de Gestió d’Ajuts Universitaris i de Recerca (AGAUR) (2016SGR924) and by the Tropical Disease Cooperative Research Network (RICET) (RD16/0027/0004). ISGlobal is a member of the Centres de Recerca de Catalunya (CERCA) Programme, Government of Catalonia (Spain).

## Conflict of interest

The funders had no role in study design, data collection and analysis, decision to publish, or preparation of the manuscript. None of the authors declares having conflicts of interest.

## Supporting Information Legends

S1 Checklist: STROBE Checklist

